# *Brainrender*: a python-based software for visualizing anatomically registered data

**DOI:** 10.1101/2020.02.23.961748

**Authors:** F. Claudi, A. L. Tyson, L. Petrucco, T.W. Margrie, R. Portugues, T. Branco

## Abstract

The recent development of high-resolution three-dimensional (3D) digital brain atlases and high-throughput brain wide imaging techniques has fueled the generation of large datasets that can be registered to a common reference frame. This registration facilitates integrating data from different sources and resolutions to assemble rich multidimensional datasets. Generating insights from these new types of datasets depends critically on the ability to easily visualize and explore the data in an interactive manner. This is, however, a challenging task. Currently available software is dedicated to single atlases, model species or data types, and generating 3D renderings that merge anatomically registered data from diverse sources requires extensive development and programming skills. To address this challenge, we have developed *brainrender*: a generic, open-source Python package for simultaneous and interactive visualization of multidimensional datasets registered to brain atlases. *Brainrender* has been designed to facilitate the creation of complex custom renderings and can be used programmatically or through a graphical user interface. It can easily render different data types in the same visualization, including user-generated data, and enables seamless use of different brain atlases using the same code base. In addition, *brainrender* generates high-quality visualizations that can be used interactively and exported as high-resolution figures and animated videos. By facilitating the visualization of anatomically registered data, *brainrender* should accelerate the analysis, interpretation, and dissemination of brain-wide multidimensional data.

## Introduction

Understanding how nervous systems generate behavior benefits from gathering multi-dimensional data from different individual animals. These data range from neural activity recordings and anatomical connectivity, to cellular and subcellular information such as morphology and gene expression profiles. These different types of data should ideally all be in register so that, for example, neural activity in one brain region can be interpreted in light of the connectivity of that region or the cell types it contains. Such registration, however, is challenging. Often it is not technically feasible to obtain multi-dimensional data in a single experiment, and registration to a common reference frame must be performed post-hoc. Even for the same experiment type, registration is necessary to allow comparisons across individual animals.

While different types of references can in principle be used, neuroanatomical location is a natural and most commonly used reference frame (Chon et al. 2019; Oh et al. 2014; Arganda-Carreras et al. 2018; Kunst et al. 2019). In recent years, several high-resolution threedimensional electronic brain atlases have been generated for model species commonly used in neuroscience (e.g.: Wang et al. 2020; Kunst et al. 2019; Arganda-Carreras et al. 2018). These atlases provide a framework for registering different types of data across macro- and microscopic scales. A key output of this process is the visualization of all datasets in register. Given the intrinsically three-dimensional (3D) geometry of brain structures and individual neurons, 3D renderings are more readily understandable and can provide more information when compared to two dimensional images. Exploring interactive 3D visualizations of the brain gives an overview of the relationship between datasets and brain regions and helps generating intuitive insights about these relationships. This is particularly important for large-scale datasets such as the ones generated by open-science projects like MouseLight (Winnubst et al. 2019) and the Allen Mouse Connectome (Oh et al. 2014). In addition, high-quality 3D visualizations facilitate the communication of experimental results registered to brain anatomy.

Generating custom 3D visualizations of atlas data requires programmatic access to the atlas. While some of the recently developed atlases provide an API (Application Programming Interface) for accessing atlas data (Wang et al. 2020; Kunst et al. 2019), rendering these data in 3D remains a demanding and time-consuming task that requires significant programming skills. Moreover, visualization of user-generated data registered onto the atlas requires an interface between the user data and the atlas data, which further requires advanced programming knowledge and extensive development. There is therefore the need for software that can simplify the process of visualizing 3D anatomical data from available atlases and from new experimental datasets.

Currently, existing software packages such as [cocoframer](https://github.com/AllenInstitute/cocoframer) (Lein et al. 2007), *[BrainMesh](https://github.com/Yaoyao-Hao/BrainMesh)* (Yaoyao-Hao 2020) and *[SHARPTRACK](https://github.com/cortex-lab/allenCCF)* (Shamash et al. 2018) provide some functionality for 3D rendering of anatomical data. These packages, however, are only compatible with a single atlas and cannot be used to render data from different atlases or different animal species. Achieving this requires adapting the existing software to the different atlases datasets or developing new dedicated software all together, at the cost of significant additional efforts, often duplicated. An important limitation of the currently available software is that it frequently does not support rendering of non-atlas data, such as data from publicly available datasets (e.g.: MouseLight) or produced by individual laboratories. This capability is essential for easily mapping newly generated data onto brain anatomy at high-resolution and produce visualizations of multidimensional datasets. More advanced software such as *[natverse](https://github.com/natverse)* (Bates et al. 2020) offers extensive data visualization and analysis functionality but currently it is mostly restricted to data obtained from the *drosophila* brain. *Simple Neurite Tracer* (Arshadi et al. 2020), an ImageJ-based software, can render neuronal morphological data from public and user-generated datasets and is compatible with several reference atlases. However, this software does not support visualization of data other than neuronal morphological reconstructions nor can it be easily adapted to work with different or new atlases beyond the ones already supported. Finally, software such as *MagellanMapper* (Young et al. 2020) can be used to visualize and analyze large 3D brain imaging datasets, but the visualization is restricted to one data item (i.e. images from one individual brain). It is therefore not possible to combine data from different sources into a single visualization. Ideally, a rendering software should work with 3D mesh data instead of 3D voxel image data to allow the creation of high-quality renderings and facilitate the integration of data from different sources.

An additional consideration is that existing software tools for programmatic neuroanatomical renderings have been developed in programming languages such as R and Matlab, and there is currently no available alternative in Python. The popularity of Python within the neuroscientific community has grown tremendously in recent years (Muller et al. 2015). Building on Python’s simple syntax and free, high-quality data processing and analysis packages, several open-source tools directly aimed at neuroscientists have been written in Python and are increasingly used (e.g. see Mathis et al. 2018; Pachitariu et al. 2017; Tyson et al. 2020). Developing a python-based software for universal generation of 3D renderings of anatomically registered data can therefore take advantage of the increasing strength and depth of the python neuroscience community for testing and further development.

For these reasons we have developed *brainrender*: an open-source python package for creating high-resolution, interactive 3D renderings of anatomically registered data. *Brainrender* is written in Python and integrated with BrainGlobe’s *AtlasAPI* (Claudi, Tyson, Petrucco et al. 2020) to interface natively with different atlases without need for modification. *Brainrender* supports the visualization of data acquired with different techniques and at different scales. Data from multiple sources can be combined in a single rendering to produce rich and informative visualizations of multi-dimensional data. *Brainrender* can also be used to create high-resolution, publication-ready images and videos (see Tyson et al. 2020; Adkins et al. 2020), as well as interactive online visualizations to facilitate the dissemination of anatomically registered data. Finally, using *brainrender* requires minimal programming skills, which should accelerate the adoption of this new software by the research community. All *brainrender* code is available at the [GitHub repository](https://github.com/brainglobe/brainrender) together with extensive online [documentation](https://docs.brainrender.info/) and examples.

## Results

### Design principles and implementation

A core design goal for *brainrender* was to generate a visualization software compatible with any reference atlas, thus providing a generic and flexible tool (Figure 1A). To achieve this goal, *brainrender* has been developed as part of the BrainGlobe’s computational neuroanatomy software suite. In particular, we integrated *brainrender* directly with BrainGlobe’s *AtlasAPI* (Claudi, Tyson, Petrucco et al. 2020). The *AtlasAPI* can download and access atlas data from several supported atlases in a unified format and new atlases can be easily adapted to work with the API. *Brainrender* uses the *AtlasAPI* to access 3D mesh data from individual brain regions as well as metadata about the hierarchal organization of the brain’s structures (Figure 1B). Thus, the same programming interface can be used to access data from any atlas (see code examples in Figure 2), including recently developed ones (e.g.: the enhanced and unified mouse brain atlas, Chon et al. 2019).

**Figure 1.**
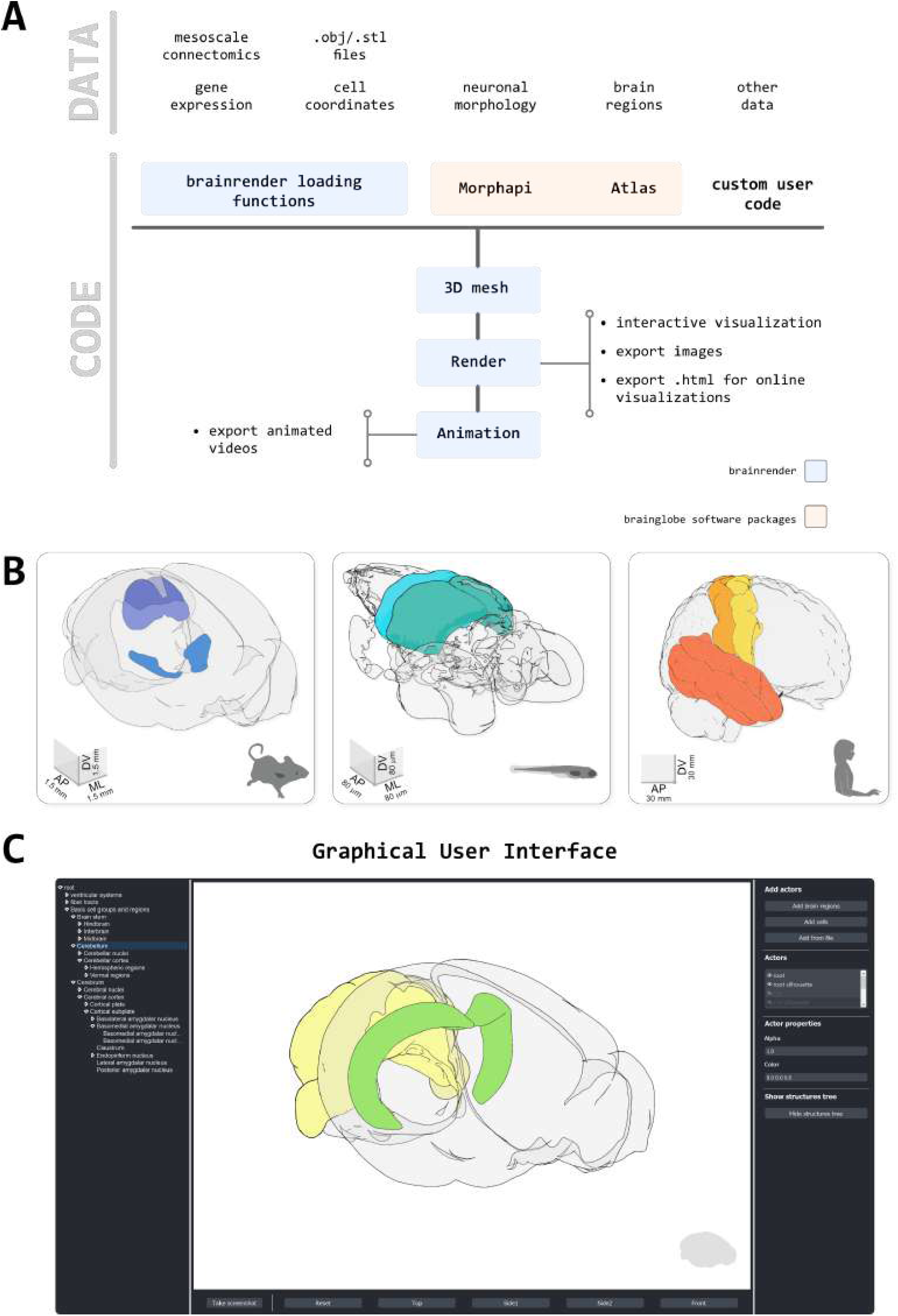
*Brainrender* design principles. **A)** Schematic illustration of how different types of data can be loaded into *brainrender* using either *brainrender*’s own functions, software packages from the BrainGlobe suite or custom Python scripts. All data loaded into *brainrender* is converted onto a unified format, which simplifies the process of visualizing data from different sources. **B)** Using *brainrender* with different atlases. Visualization of brain atlas data from three different atlases using *brainrender*. Left, Allen atlas of the mouse brain showing the superficial (SCs) and motor (SCm) subdivisions of the superior colliculus and the Zona Incerta (data from Wang et al. 2020). Middle, visualization of the cerebellum and tectum in the larval zebrafish brain (data from Kunst et al. 2019). Right, visualization of the precentral gyrus, postcentral gyrus and temporal lobe of the human brain (data from Ding et al. 2016). **C)** The *brainrender* GUI. Mouse, human and zebrafish larvae drawings from scidraw.io (doi.org/10.5281/zenodo.3925991, doi.org/10.5281/zenodo.3926189, doi.org/10.5281/zenodo.3926123)

**Figure 2.**
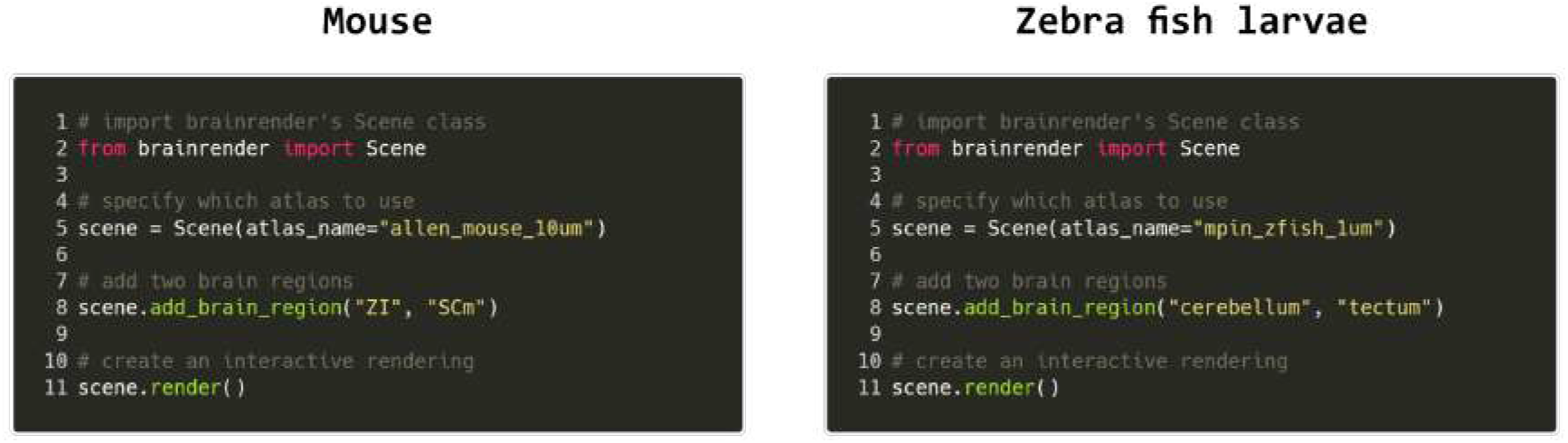
Code examples. Example python code for visualizing brain regions in the mouse and larval zebrafish brains. The same commands can be used for both atlases and switching between atlases can be done by simply specifying which atlas to use when creating the visualization.

The second major design principle was to enable rendering of any data type that can be registered to a reference atlas, either from publicly available datasets or from individual laboratories. To achieve this, all data loaded in *brainrender* is represented as 3D mesh information, which enables the use of state-of-the-art rendering tools (Hanwell et al. 2015; Musy, Dalmasso, and Sullivan 2019). Converting data into a 3D mesh data format is a non-trivial and time-consuming task that requires significant programming expertise. To facilitate this process, *brainrender* provides functionality for easily loading and visualizing commonly used data types, such as the location of labelled cells or 3D mesh data from .obj and .stl files. In addition, *brainrender* can visualize data produced with any analysis software from the BrainGlobe suite, including *cellfinder* (Tyson et al. 2020) and *brainreg* (Tyson, Rousseau, and Margrie 2020). The existing loading functionality can be easily expanded to support user-specific needs by directly plugging in custom user code into the *brainrender* interface (Figure 1A).

One of the goals of *brainrender* is to facilitate the creation of high-resolution images, animated videos and interactive online visualizations from any anatomically registered data. *Brainrender* uses *vedo* as the rendering engine (Musy, Dalmasso, and Sullivan 2019), a state-of-the-art tool that enables fast, high-quality rendering with minimal hardware requirements (e.g.: no dedicated GPU is needed). Animated videos and online visualizations can be produced with a few lines of code in *brainrender*. Several options are provided for easily customizing the appearance of rendered objects, thus enabling high-quality, rich data visualizations that combine multiple data sources.

Finally, we aimed for *brainrender* to empower scientists with little, or no programming experience to generate advanced visualizations of their anatomically registered data. To make *brainrender* as user-friendly as possible we have produced extensive [documentation](https://docs.brainrender.info/), tutorials and examples for installing and using the software. We have also developed a Graphic User Interface (GUI) to access most of *brainrender*’s core functionality. This GUI can be used to perform actions such as rendering of brain regions and labelled cells (e.g.: from *cellfinder*) and creating images of the rendered data, without writing custom python code (Figure 1C).

### Visualizing brain regions and other structures

A key element of any neuroanatomical visualization is the rendering of the entire outline of the brain as well as the borders of brain regions of interest. In *brainrender* this can easily be achieved by specifying which brain regions to include in the rendering. The software will then use Brain-Globe’s *AtlasAPI* to load the 3D data and subsequently renders them (Figure 1B).

*brainrender* can also render brain areas defined by factors other than anatomical location, such as gene expression levels or functional properties. These can be loaded either directly as 3D mesh data after processing with dedicated software (e.g. Tyson, Rousseau, and Margrie 2020; Song et al. 2020; Jin et al. 2019)(Figure 3A), or as 3D volumetric data (Figure 3E). For the latter, *brainrender* takes care of the conversion of voxels into a 3D mesh for rendering. Furthermore, custom 3D meshes can be created to visualize different types of data. For example, *brainrender* can import JSON files with tractography connectivity data and create ‘streamlines’ to visualize efferent projections from a brain region of interested (Figure 3B).

**Figure 3.**
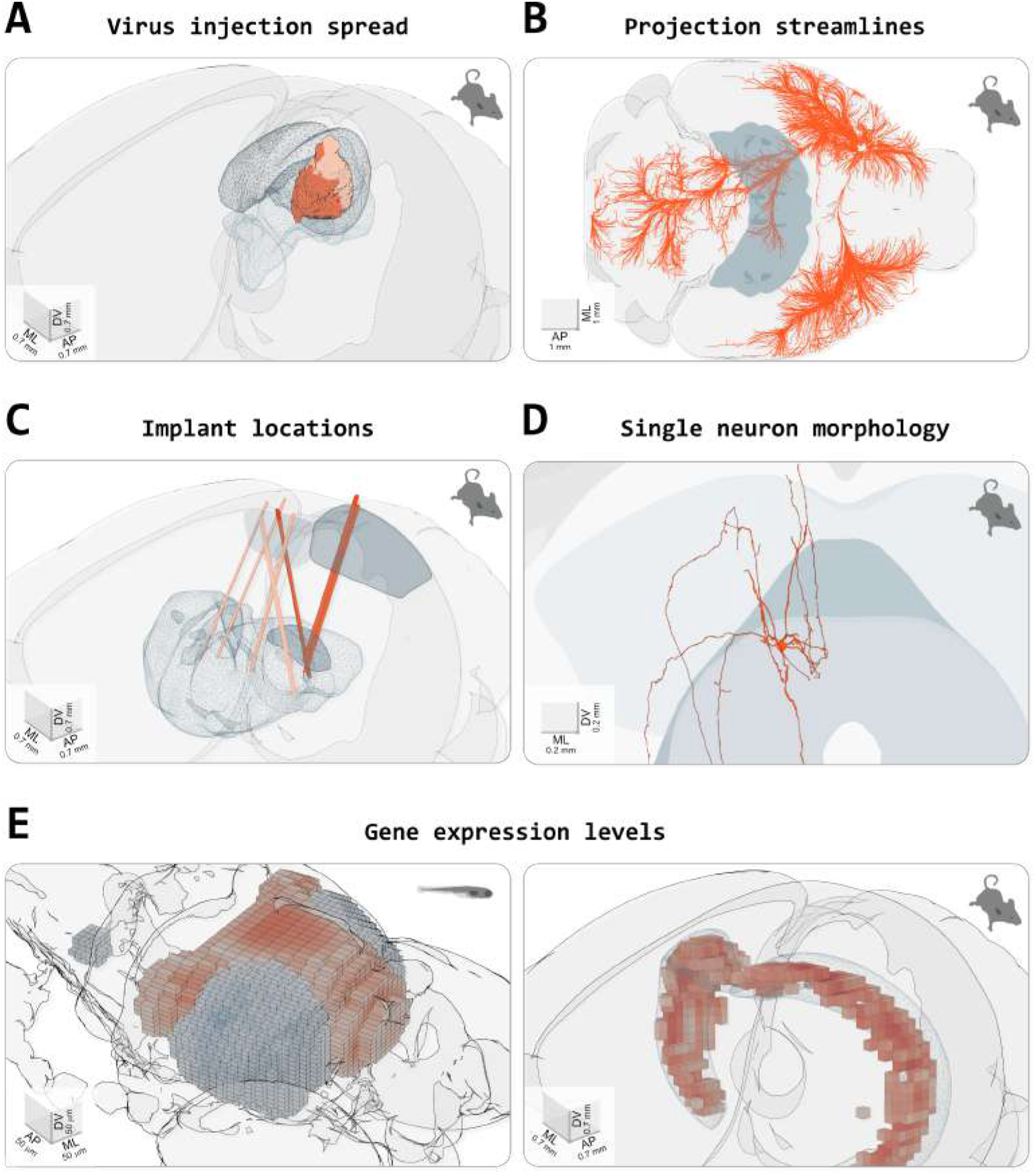
Visualizing different types of data in *brainrender*. **A)** Spread of fluorescence labelling following viral injection of AAV2-CRE-eGPF in the superior colliculus of two FLEX-TdTomato mice. 3D objects showing the injection sites were created using custom python scripts following acquisition of a 3D image of the entire brain with serial 2-photon tomography and registration of the image data to the atlas’ template (with *brainreg*, Tyson, Rousseau, and Margrie 2020). **B)** Streamlines visualization of efferent projections from the mouse primary motor cortex following injection of an anterogradely transported virus expressing fluorescent proteins (original data from Oh et al. 2014, downloaded from [Neuroinformatics NL](https://neuroinformatics.nl/drupal) with *brainrender*). **C)** Visualization of the location of several implanted neuropixel probes from multiple mice (data from Steinmetz et al. 2019). Dark salmon colored tracks show probes going through both primary/anterior visual cortex (VISp/VISa) and the dorsal lateral geniculate nucleus of the thalamus. **D)** Single periaqueductal gray (PAG) neuron. The PAG and superior colliculus are also shown. The neuron’s morphology was reconstructed by targeting the expression of fluorescent proteins in excitatory neurons in the PAG via an intersectional viral strategy, followed by imaging of cleared tissue and manual reconstruction of the neuron’s morphology with Vaa3D software. Data were registered to the Allen atlas with *SHARPTRACK* (Shamash et al. 2018). The 3D data was saved as a .stl file and loaded directly into *brainrender*. **E)** Gene expression data. Left, expression of genes ‘brn3c’ and ‘nk1688CGt’ in the tectum of the larval zebrafish brain (gene expression data from fishatlas.neuro.mpg.de, 3D objects created with custom python scripts). Right, expression of gene ‘Gpr161’ in the mouse hippocampus (gene expression data from Wang et al. 2020, downloaded with *brainrender*. 3D objects created with *brainrender*). Colored voxels show voxels with high gene expressions. The CA1 field of the hippocampus is also shown.

*Brainrender* also simplifies visualizing the location of devices implanted in the brain for neural activity recordings or manipulations, such as electrodes or optical fibers. Post-hoc histological images taken to confirm the correct placement of the device can be registered to a reference atlas using appropriate software and the registered data can be imported into *brainrender* (Figure 3C). This type of visualization greatly facilitates cross-animal comparisons and helps data interpretation within and across research groups.

Finally, *brainrender* can be used to visualize any object represented by the most commonly used file formats for three-dimensional design (e.g.: .obj, .stl), thus ensuring that *brainrender* can flexibly adapt to the visualization needs of the user (Figure 3D).

### Individual neurons and mesoscale connectomics

Recent advances in large field of view and whole-brain imaging allow the generation of brain-wide data at single neuron resolution. Having a platform for visualizing these datasets with ease is critical for exploratory data analyses. Several open source software packages are available for and registering large amounts of such imaging data (Tyson et al. 2020; Fürth et al. 2018; Goubran et al. 2019; Renier et al. 2016), and automatically identify labelled cells (e.g.: expressing fluorescent proteins). This processing step outputs a table of coordinates for a set of labelled cells, which can be directly imported into *brainrender* to visualize a wealth of anatomical data at cellular resolution (Figure 4A).

**Figure 4.**
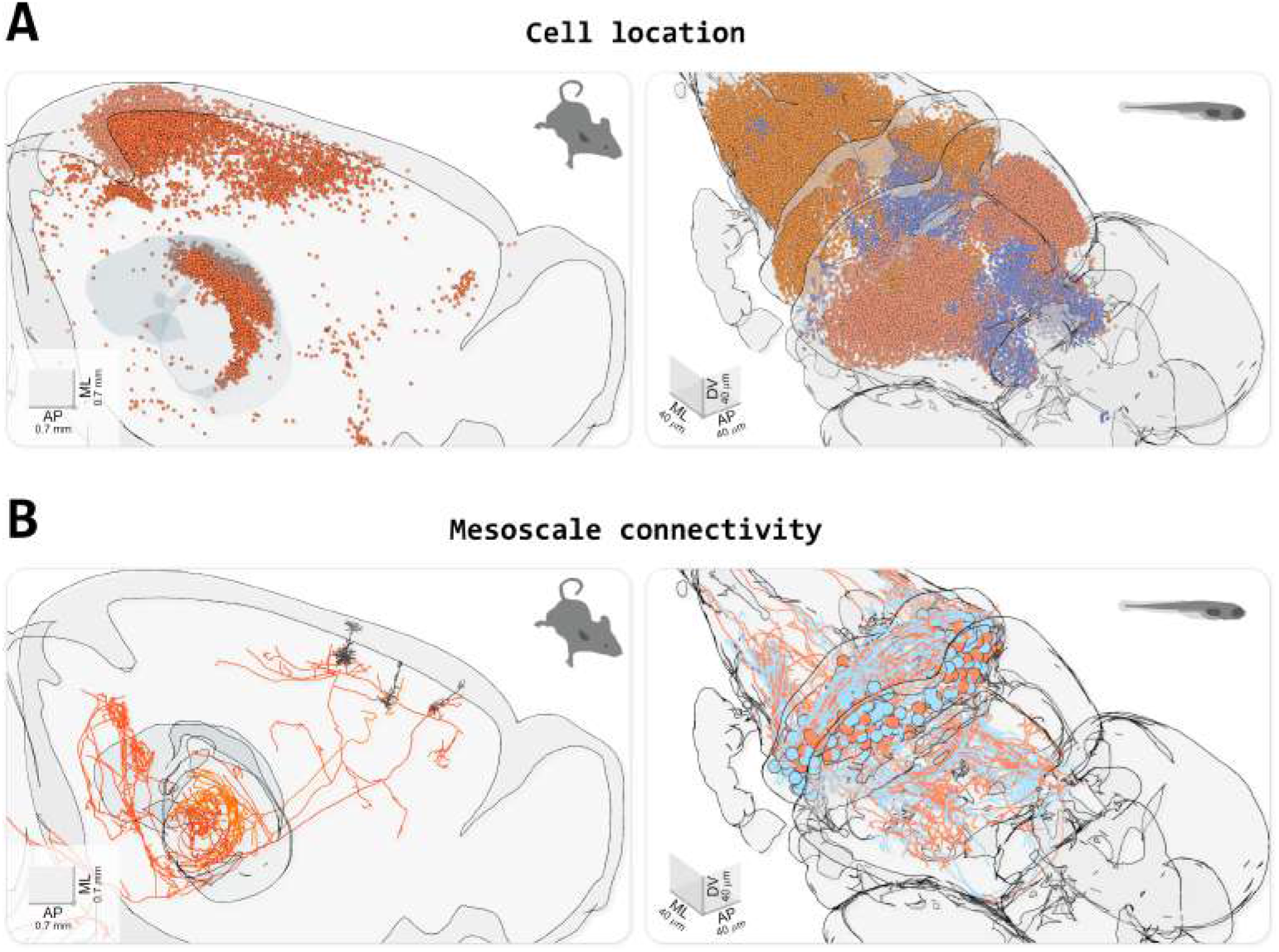
Visualizing cell location and morphological data. **A)** Visualizing the location of labelled cells. Left, visualization of fluorescently labelled cells identified using *cellfinder* (data from Tyson et al. 2020). Right, visualization of functionally defined clusters of regions of interest in the brain of a zebrafish larvae during a visuomotor task. (data from Markov et al. 2020). **B)** Visualizing neuronal morphology data. Left. three secondary motor cortex neurons projecting to the thalamus (data from Winnubst et al. 2019, downloaded with *morphapi* from neuromorpho.org). Right, morphology of cerebellar neurons in larval zebrafish (data from Kunst et al. 2019, downloaded with *morphapi*). In the left panel of A) and B), the brain’s outline was sliced along the midline to expose the data.

Beyond the location of cell bodies, visualizing the entire dendritic and axonal arbors of single neurons registered to a reference atlas is important for understanding the distribution of neuronal signals across the brain. Single cell morphologies are often complex threedimensional structures and therefore poorly represented in two-dimensional images. Generating three-dimensional interactive renderings is thus important to facilitate the exploration of this type of data. *brainrender* can be used to parse and render .swc files containing morphological data and it is fully integrated with *morphapi*, a software for downloading morphological data from publicly available datasets (e.g.: from neuromorpho.org) (Figure 4B).

### Producing figures, videos and interactive visualizations with *brainrender*

A core goal of *brainrender* is to facilitate the production of high-quality images, videos and interactive visualizations of anatomical data. *brainrender* leverages the functionality provided by *vedo* (Musy, Dalmasso, and Sullivan 2019) to create images directly from the rendered scene. Renderings can also be exported to HTML files to create interactive visualizations that can be hosted online. Finally, functionality is provided to easily export videos from rendered scenes. Animated videos can be created by specifying parameters (e.g.: the position of the camera or the transparency of a mesh) at selected keyframes. *brainrender* then creates a video by animating the rendering between the keyframes. This approach facilitates the creation of videos while retaining the flexibility necessary to produce richly animated sequences ([Videos 1-4](https://figshare.com/articles/media/Claudi_et_al_2020_-_Brainrender_videos/13359785)). All example figures and videos in this article were generated directly in *brainrender*, with no further editing.

## Conclusions

In this article we have presented *brainrender*, a python software for creating three-dimensional renderings of anatomically registered data. *brainrender* builds on BrainGlobe’s *AtlasAPI* to provide a user-friendly, yet powerful visualization tool. The integration with the BrainGlobe ecosystem enables seamless use of different brain atlases using the same code base. This feature allows the production of generic code that can be shared and re-used for different purposes, thereby speeding up visualization developments. The integration with BrainGlobe also allows direct visualization of data generated with tools such as *brainreg* and *cellfinder*, as well of data downloaded from publicly available datasets through *morphapi*. The interoperability of multiple software packages dedicated to different tasks facilitates the development of analysis pipelines (Bates et al. 2020) and it is one of the core principles motivating the development of *brainrender*. By making the rendering processes as easy as possible, including access to most of the functionality through a GUI, we have tried to facilitate as much as possible the adoption of *brainrender* for generating high quality data visualizations of anatomically registered data.

We have demonstrated the use of *brainrender* for rendering different types of data, including brain-wide gene expression profiles, mesoscale connectivity patters and single neuron morphologies. These include user-generated data and data from large scale projects such as Mouse-Light and the Allen Institute’s Mouse Connectome projects. While we have aimed to make the visualization process as easy as possible, this is not at the cost of flexibility. Rendering different types of data and advanced custom visualizations can be achieved by plugging in custom code directly into the *brainrender* engine. Our software is accompanied by extensive online documentation, tutorials and examples to facilitate adoption and achieve the goal of providing a useful, open-source tool for the community. In addition, the code has been written following best practices (e.g.: thorough testing and documentation) and therefore provide a solid base for future developments.

### Limitations and future directions

While we have designed *brainrender* usage to require minimal programming expertise, installing python and *brainrender* may still prove challenging for some users. In the future, we aim to make *brainrender* a stand-alone application that can be simply downloaded and locally installed.

In addition to images and videos, *brainrender* can be used to export renderings as HTML files and generate online 3D interactive renderings. Currently, however, embedding renderings into a web page remains far from a trivial task. Further developments on this front should make it possible to easily host interactive renderings online, therefore improving how anatomically registered data are disseminated both in scientific publications and other media.

## Methods

*Brainrender* is written in Python 3 and depends on standard python packages (Harris et al. 2020) and on *vedo* (Musy, Dalmasso, and Sullivan 2019) and BrainGlobe’s *AtlasAPI* (Claudi, Tyson, Petrucco et al. 2020). Extensive documentation on how to install and use *brainrender* can be found at docs.brainrender.info and we provide here a only brief overview of the workflow in *brainrender*. The GitHub repository also contains detailed examples of Python scripts and Jupyter notebooks. All *brainrender*’s code is open-source and has been deposited in full in the [GitHub repository](https://github.com/brainglobe/brainrender) and at PyPi (a repository of Python software) under a permissive BSD 3-Clause license. We welcome any user to download and inspect the source code, modify it as needed or contribute to *brainrender*’s development directly

### Brainrender’s workflow

The central element of any visualization produced by *brainrender* is the Scene. A Scene controls which elements (Actors) are visualized and coordinates the rendering, the position of the camera’s point of view, the generation of screenshots and animations from the rendered scene and other important actions.

Actors can be added to the scene in several ways. When loading data directly from a file with 3D mesh information (e.g.: .obj) an Actor is generated automatically to represent the mesh in the rendering. When rendering data from other sources (e.g.: from a .swc file with neuronal morphology or from a table of coordinates of labelled cells), dedicated functions in *brainrender* parse the input data and generate the corresponding Actors. Actors in *brainrender* have properties, such as color and transparency, that can be used to specify the appearance of a rendered actor accordingly to the user’s aesthetic preferences. *Brainrender*’s Scene and Actor functionality use *vedo* as the rendering engine ([GitHub repository](https://github.com/marcomusy/vedo); Musy, Dalmasso, and Sullivan 2019).

In addition to data loaded from external files, *brainrender* can directly load atlas data containing, for example, the 3D meshes of individual brain regions. This is done via BrainGlobe’s *AtlasAPI* to allow the same programming interface in *brainrender* to visualize data from any atlas supported by the *AtlasAPI. Brainrender* also provides additional functionality to interface with data available from projects that are part of the Allen Institute Mouse Atlas and Mouse Connectome projects (Wang et al. 2020; Oh et al. 2014). These projects provide an SDK (Software Development Kit) to directly download data from their database and *brainrender* provides a simple interface for downloading gene-expression and connectomics (streamlines) data. All atlas and connectomics data downloaded by *brainrender* can be loaded directly into a Scene as Actors.

Visualizing morphological data with reconstructions of individual neurons can be done by loading these type of data directly from .swc files, or by downloading them in Python using *morphapi* - software from the BrainGlobe suite that provides a simple and unified interface with several databases of neurons morphologies (e. g. neuromorpho.org). Data downloaded with *morphapi* can be loaded directly into a *brainrender* scene for visualization.

### Example code

As a demonstration of how easily renderings can be created in *brainrender*, the following Python code illustrates how to create a Scene and add Actors by loading 3D data from an .obj file and then adding brain regions to the visualization (Figure 5).

**Figure 5.**
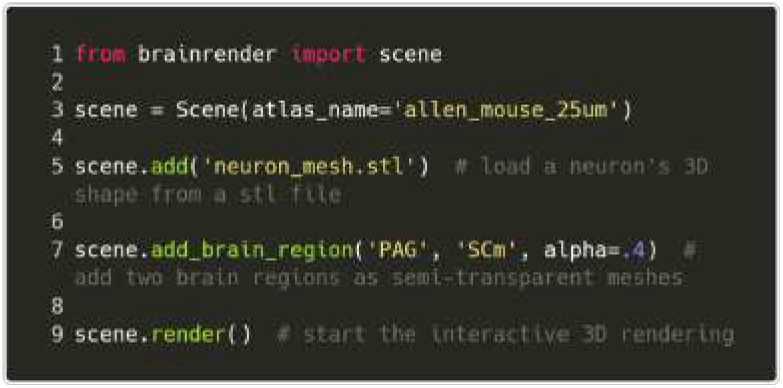
Example code.

The code used to generate the figures and videos in this article is made freely available at a dedicated [GitHub repository](https://github.com/brainglobe/brainrender_paper).

## Acknowledgements

We thank Yu Lin Tan for sharing the single neuron morphology shown in Figure 3D. We thank scidraw.io for the illustrations of a human, mouse and zebrafish used in Figures 1, 2 and 3. This work was supported by grants from the GatsbyCharitable Foundation (GAT3361, T.W.M. and T.B.), Wellcome Trust (090843/F/09/Z,T.W.M. and T.B.; 214333/Z/18/Z, T.W.M.; 214352/Z/18/Z, T.B.) and by the DeutscheForschungsgemeinschaft (DFG, German Research Foundation) under Germany’s Excellence Strategy within the framework of the Munich Cluster for Systems Neurology (EXC 2145 SyN-ergy — ID 390857198).

